# Spatial variation of the native colon microbiota in healthy adults

**DOI:** 10.1101/189886

**Authors:** Kaitlin J. Flynn, Mack T. Ruffin, D. Kim Turgeon, Patrick D. Schloss

## Abstract

The microbiome has been implicated in the development of colorectal cancer (CRC) and inflammatory bowel diseases (IBD). The specific traits of these diseases vary along the axis of the digestive tract. Further, variation in the structure of the gut microbiota has been associated with both diseases. Here we profiled the microbiota of the healthy proximal and distal mucosa and lumen to better understand how bacterial populations vary along the colon. We used a two-colonoscope approach to sample proximal and distal mucosal and luminal contents from the colons of 20 healthy subjects that had not undergone any bowel preparation procedure. The biopsies and home-collected stool were subjected to 16S rRNA gene sequencing and Random Forest classification models were built using taxa abundance and location to identify microbiota specific to each site. The right mucosa and lumen had the most similar community structures of the five sites we considered from each subject. The distal mucosa had higher relative abundance of *Finegoldia, Murdochiella, Peptoniphilus, Porphyromonas* and *Anaerococcus*. The proximal mucosa had more of the genera *Enterobacteriaceae, Bacteroides* and *Pseudomonas*. The classification model performed well when classifying mucosal samples into proximal or distal sides (AUC = 0.808). Separating proximal and distal luminal samples proved more challenging (AUC = 0.599) and specific microbiota that differentiated the two were hard to identify. By sampling the unprepped colon, we identified distinct bacterial populations native to the proximal and distal sides. Further investigation of these bacteria may elucidate if and how these groups contribute to different disease processes on their respective sides of the colon.

## Introduction

The human colon is an ecosystem comprised of numerous microenvironments that select for different microbiota. Concentrations of oxygen, water, and anti-microbial peptides change along the gut axis and influence which microbiota reside in each location. Microenvironments differ not only longitudinally along the colon, but also radially from the epithelium to mucosa to intestinal lumen, offering several sites for different microbial communities to flourish. The identity of these specific microbiota and communities are important for understanding the etiology of complex diseases such as Colorectal Cancer (CRC) and Inflammatory Bowel Disease (IBD). CRC and IBD can be preceded or accelerated by perturbations of the structure of the gut microbiota (1–3). The manifestations of these diseases are known to vary based upon the location in which they occur. For instance, CRC that arises in the distal (left) colon are of hindgut origin and tend to have large chromosomal alterations indicative of chromosomal instability (1). In contrast, CRC arising in the proximal (right) colon are of midgut origin and tend to be sessile and microsatellite instable (MSI with BRAF and KRAS mutations) (1). In addition to the environmental gradients within the colon, the distal and proximal sides of the colon differ in the amount of inflammation present and the genomic instability of precancerous cells, respectively (1,4,5). In IBD patients, disease occurring in the distal colon extending proximally is usually indicative of ulcerative colitis (UC), whereas Crohn’s disease (CD) can occur anywhere along the GI tract, most commonly in the ileum and the cecum (2). UC presents as continuous disease with only mucosal involvement, where as CD has skip lesions and full thickness involvement that may cause abscesses, strictures and fistulas (2). Thus, given the varied physiology of the proximal-distal axis of the colon and known differences in disease patterns at these sites, symbiotic microbiota and their metabolites likely vary as well, and may influence the heterogeneous disease prognoses of IBD and CRC. Because CRC can be a long-term complication of IBD, the distribution of microbiota is important to understanding the pathophysiology of both diseases.

Several recent findings have shown that development and progression of IBD or CRC can be attributed to specific molecular events as a result of interactions between the gut microbiota and human host (1,3,6). For instance, comparison of the bacteria present on CRC tumors with those found on nearby healthy tissue has identified specific species that are tumor-associated (7). Specific bacteria have also been identified in fecal samples of patients with varying stages of colon tumorigenesis (8,9). These species include the oral pathogens *Fusobacterium nucleatum* and *Porphyromonas asacharolytica. F. nucleatum* has also been found to be elevated in the stool and biopsies of patients with IBD as compared to healthy controls (10,11). Furthermore, studies of *F. nucleatum* isolated from mucosal biopsies showed that more invasive *F. nucleatum* positively correlates with IBD disease level (10). Like many intestinal pathogens, the bacteria appear to have a high-impact despite being lowly-abundant in the community (2). The physiology of these rare taxa may contribute to the colonic disease state. These studies often examined only shed human stool or the small intestine, preventing fine-resolution analysis of paired samples from the proximal and distal sides of the colon. Similarly, comparisons of on- or off-tumor/lesion bacteria rarely have matched tissue from the other side of the colon from the same, disease-baring patient, limiting what conclusions can be drawn about the colonic microbiome overall, let alone at that specific site (12). Due to these limitations, the contribution of the gut microbiota to CRC and IBD disease location in the colon is largely undefined. Characterizing these communities in healthy individuals could provide needed insight into disease etiology, including how the disruption of the healthy community could promote the initiation or proliferation of the distinct proximal and distal CRC tumors or IBD flares.

The few existing profiles of the microbial spatial variation of the colon have been limited by sample collection methods. The majority of human gut microbiome studies have been performed on whole shed feces or on samples collected during colonoscopy or surgery (5). While invasive methods allow investigators to acquire samples from inside the human colon, typically these procedures are preceded by the use of bowel preparation methods such as the consumption of laxatives to cleanse the bowel. Bowel preparation is essential for detecting cancerous or precancerous lesions in the colon, but complicates microbiome profiling as the chemicals strip the bowel of contents and disrupt the mucosal layer (13,14). As such, what little information we do have about the spatial distribution of the microbiota in the proximal and distal colon is confounded by the bowel preparation procedure.

Here we address the limitations of previous studies and identify the microbes specific to the lumen and mucosa of the proximal and distal healthy human colon. We used an unprepared colonoscopy technique to sample the natural community of each location of the gut without prior disruption of the native bacteria in 20 healthy volunteers. To address the inherent inter-individual variation in microbiota, we used a machine-learning classification algorithm trained on curated 16S rRNA sequencing reads to identify the microbiota that were specific to each location. We found that our classification models were able to separate mucosal and luminal samples as well as differentiate between sides of the colon based on populations of particular microbiota. By identifying the distinguishing microbiota we are poised to ask if and how the presence or disruption of the microbiota at each site contribute to the development of the tumor subtypes of CRC in the proximal and distal human colon.

## Methods

### Human subjects

The procedures in this study and consent were approved by the Institutional Review Board at the University of Michigan Health System with protocol number HUM00082721. Subjects were recruited using the online recruitment platform and were pre-screened prior to enrollment in the study. Exclusion criteria included: use of asprin or NSAIDs within 7 days, use of antibiotics within 3 months, current use of anticoagulants, known allergies to Fentanyl, Versed and Benadryl, prior history of colon disease, diabetes, abdominal surgery, respiratory, liver, kidney or brain impairments, undergoing current chemotherapy or radiation treatment and subjects that were pregnant or trying to conceive. 20 subjects that met the criteria were selected and provided signed informed consent prior to the procedure. There were 13 female and 7 male subjects ranging in age from 25 to 64. 18 of the 20 subjects had not used antibiotics within a year prior to the collection date and 2 had not used antibiotics within 6 months. None of the subjects had medical conditions requiring frequent or extended antibiotic use.

### Sample collection

At a baseline visit, subjects gave consent and were given a home collection stool kit (Zymo). One to seven days prior to the scheduled colonoscopy, subjects collected whole stool at home and shipped the samples to a research coordinator on ice. Notably, subjects did not undergo any bowel preparation method prior to sampling. On the procedure day, subjects reported to the Michigan Clinical Research Unit at the University of Michigan Health System. Subjects were consciously sedated using Fentanyl, Versed and/or Benadryl as appropriate. A flexible sigmoidoscope was first inserted about 25cm into the colon and jumbo biopsy forceps used to collect the luminal contents. Two luminal samples were collected and the contents immediately deposited into RNAlater (Fisher) and flash-frozen in liquid nitrogen. The forceps were withdrawn and new biopsy forceps were used to collect mucosal biopsies on sections of the colon that were pink and free of stool matter. Three mucosal biopsies were collected and flash-frozen in RNAlater. These samples comprised the distal colon samples. The sigmoidoscope was then withdrawn and a pediatric colonoscope was inserted to reach the proximal colon. The proximal samples were taken from the ascending colon proximal to the hepatic flexure at 75-120cm depending on the subject. Samples were then collected in the same manner as was done in the distal colon and the colonoscope withdrawn. All samples were stored at −80°C.

### Sample processing, sequencing and analysis

DNA extraction was performed using the PowerMicrobiome DNA/RNA Isolation Kit (MO BIO Laboratories). For tissue biopsies, Bond-Breaker TCEP solution (Fisher) and 2.8mm ceramic beads (MO BIO Laboratories) were added to the bead beating step to enhance DNA recovery from mucosal samples. The resulting DNA was normalized to equal concentrations across all samples and used as template for amplification of the V4 region of the 16S rRNA gene and fragments were sequenced on an Illumina MiSeq as previously described (15). Sequences were curated using the mothur software as described previously (16). The sequences were assigned a taxonomic classification using a naive Bayesian classifier trained using a 16S rRNA gene training set from the Ribosomal Database Project (RDP) (17) and clustered into operational taxonomic units (OTUs) based on a 97% similarity cutoff. Sequencing and analysis of a mock community revealed the error rate to be 0.018%. Samples were rarefied to 4231 sequences per sample in order to reduce the effects of uneven sampling bias.

Diversity analysis was performed using the Simpson diversity calculator and *θ*_YC_ calculator metrics in mothur version 1.39.5 (16). *θ*_YC_ distances were calculated to determine the dissimilarity between two samples. Random Forest classification models were built using the AUCRF R package using a leave-one-subject out approach (18). The Random Forest models were built using the full, non-rarefied, dataset as input. For each model the data were split into a 19-subject training set and a 1-subject test set. The model was built and cross-validated using 10-fold k cross-validation (AUCRFcv) on the training set to estimate the prediction error of the model. The resultant model was then used to predict the outcome the left-out subject. This process was repeated iteratively for all subjects and results plotted as Reciever Operator Characteristic curves using the pROC R package (19). Resultant models were used to identify the OTUs that were most important for classifying each location. Species-level information for sequences of interest was obtained by aligning the sequences to the GenBank nucleotide databse using blastn. The species name was only used if the identity score was ≥ 99% over the full-length of the contig and matched a single reference.

### Statistical analysis

Differences in community membership at the phyla level were tested using the analysis of molecular variance (AMOVA) metric in mothur. Differences in *θ*_YC_ distances by location were tested using the Wilcoxon rank-sum test adjusted for multiple comparisons using the Benjamini-Hochberg procedure.

### Data availability

16S rRNA gene sequence reads and experiment metadata are available on the NCBI Sequence Read Archive (SRA) with accession number SRP124947 and PRJNA418115. A reproducible data analysis pipeline can be found at https://github.com/SchlossLab/Flynn_LRColon_CancPrevRes_2017.

## Results

### Microbial membership and diversity of the proximal and distal colon

Luminal and mucosal samples were collected from the proximal and distal colon of 20 healthy individuals who had not undergone bowel preparation (Fig. 1). Subjects also collected stool at home one week prior to the procedure. To characterize the bacterial communities present at these sites, 16S rRNA gene sequencing was performed on DNA extracted from each sample. As expected, each site was primarily dominated by *Firmicutes* and *Bacteriodetes* (Fig. 2A) (20). Samples had varying levels of diversity at each site, irrespective of the individual (Fig. 2B). For example, the proximal mucosa was more diverse than the distal for some individuals while the opposite was true for others. Therefore we could not identify a clear pattern of changes in microbial diversity along the gut axis.

**Figure 1.**
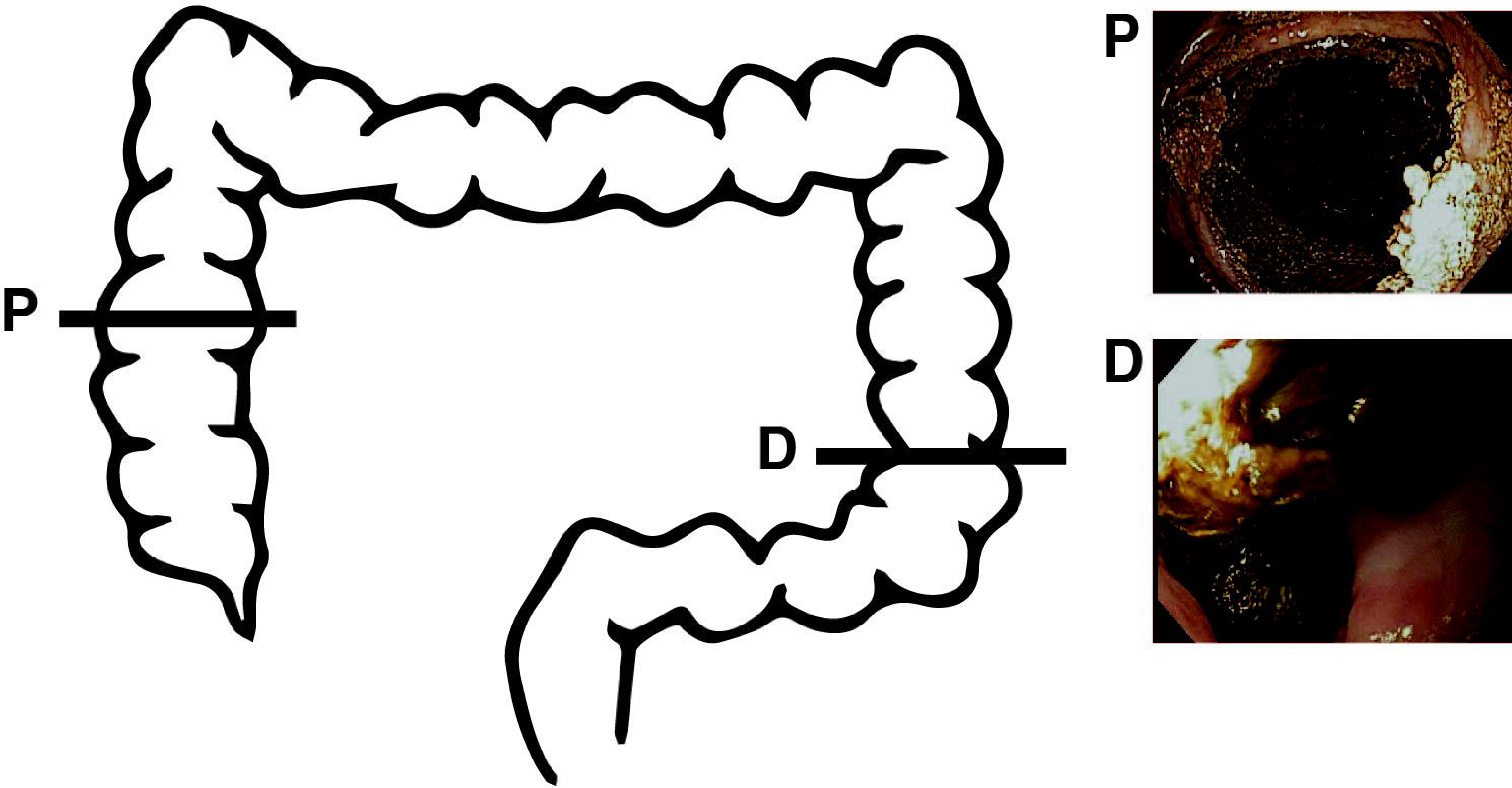
Sampling strategy. A flexible sigmoidoscope was used to sample the distal colonic luminal contents and mucosa. The scope was inserted ∼ 25cm into the subject and biopsy forceps were used to sample the luminal contents (D, inset). A separate set of biopsy forceps was used to sample the distal mucosa (D, inset). The sigmoidoscope was removed. A pediatric colonoscope was inserted and used to access the proximal colon (P, inset). Biopsies were taken of the proximal luminal contents and mucosa as described. One week prior to the procedure stool was collected at home and sent into the laboratory. Representative images from one individual are shown.

**Figure 2.**
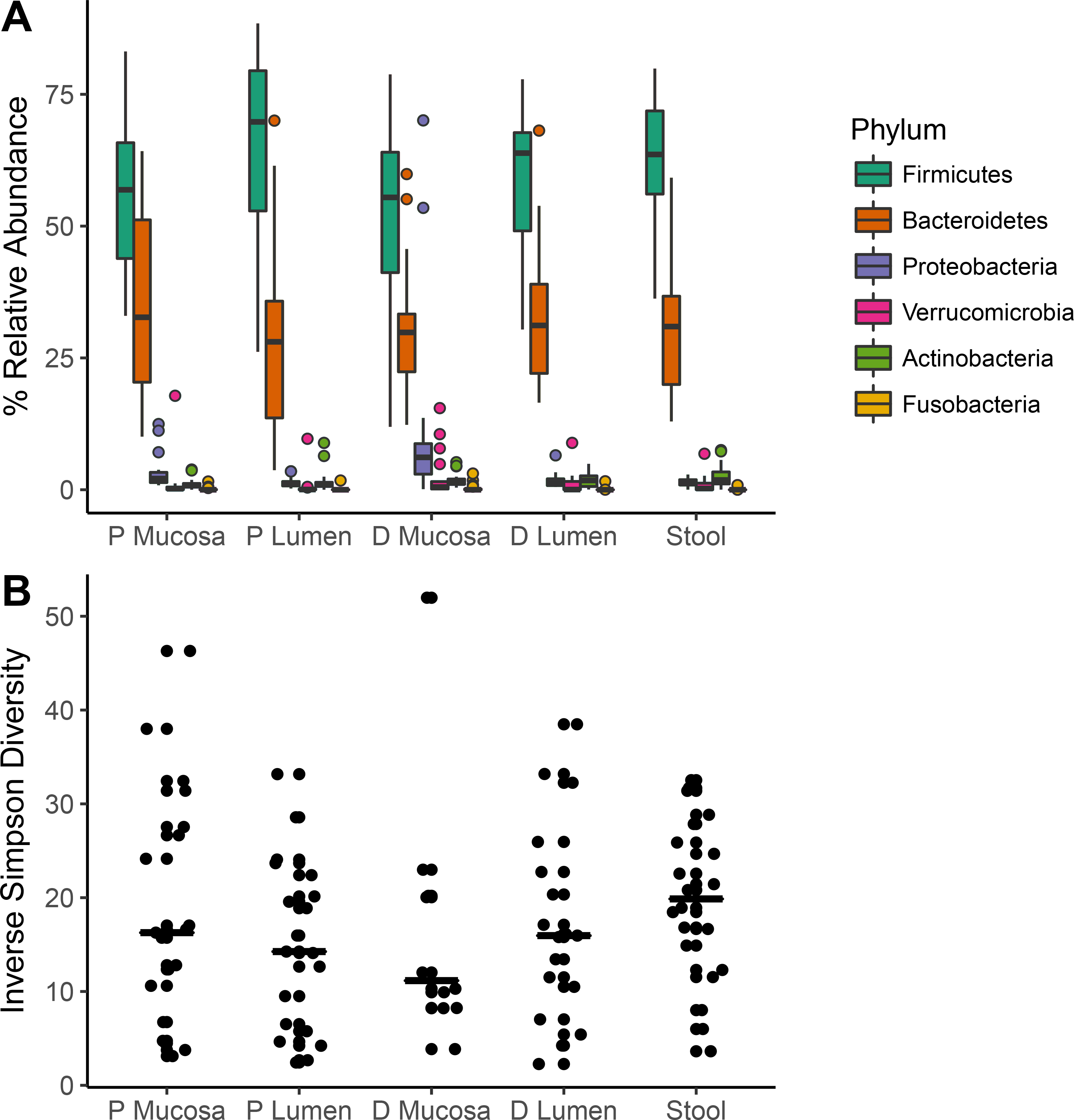
Phylum-level relative abundance and diversity in the proximal and distal human colon. A) Relative abundance of the top five bacterial phyla in each sampling site. Each box represents the median and interquartile range. B) Simpson diversity of the microbial communities at each location. The horizontal lines represent the median values.

To compare similarity between the proximal and distal sides and within the lumen and mucosa, we compared the community structure of these sites based on the relative abundances of individual Operational Taxonomic Units (OTUs). Across all subjects we observed wide variation when comparing sample locations (Fig. 3A). Those ranges did not follow a clear pattern on an individual basis. However, when comparing median dissimilarity between the communities found in the proximal lumen and mucosa, the proximal and distal lumen, the proximal and distal mucosa, and the distal lumen and mucosa, we found that the proximal lumen and mucosa were most similar to each other than to the other samples (P < 0.005, Wilcoxon, BH adjustment).

**Figure 3.**
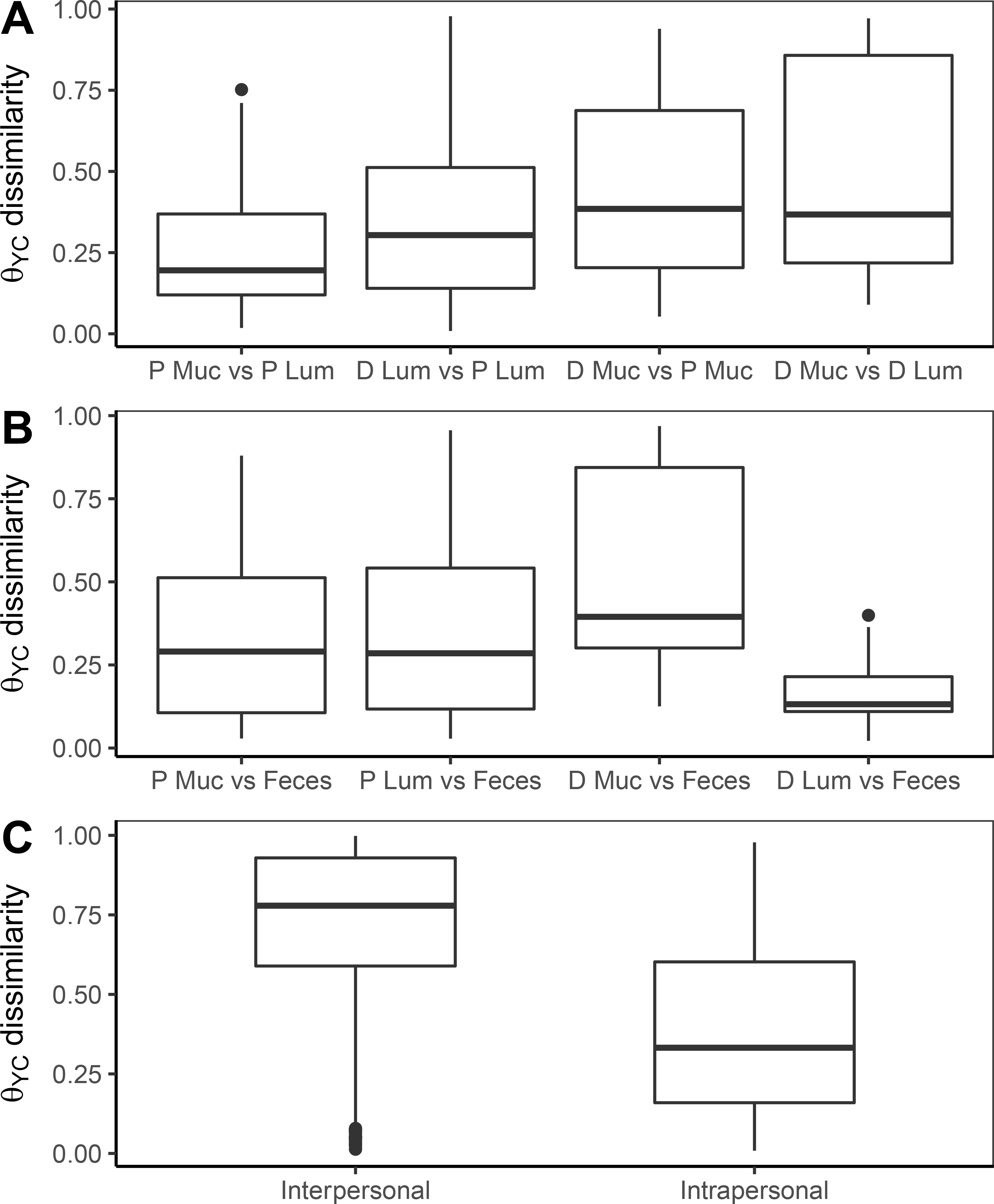
Comparison of microbial community structure between sites of the gut. *θ*_YC_ distances are shown to indicate the interpersonal dissimilarities between two sites - each point represents one individual. In (A), comparisons of the proximal and distal mucosal and lumen are shown. In (B), comparisons of each site to the exit stool are shown. In (C), comparisons of samples from all subjects to each other (interpersonal) or within one subject (intrapersonal) are shown.

### Fecal samples resemble luminal samples from the distal colon

Next, we compared the luminal and mucosal samples to the fecal sample of each subject. Amidst the large inter-subject variation, we did identify significantly less dissimilarity between the distal luminal sample and the feces (Fig. 3B, P < 0.05, Wilcoxon, BH adjustment). Furthermore, there was an even larger difference in the communities found in the distal mucosa compared to the fecal communities, indicating that the mucosa is as different from the stool as compared to lumen (P < 0.0005, Wilcoxon, BH adjustment). These results suggest that the contents of the distal lumen were most representative of the subjects’ feces, and the mucosal microbiota are distinct from the fecal and luminal communities.

### Interpersonal community variation is greater than the variation between sites

To determine what factors may have driven the differences seen among the samples, we compared the community dissimilarity between samples from all subjects (interpersonal) versus samples from within one subject (intrapersonal). We found that samples from one individual were far more similar to each other than to matched samples from the other subjects (Fig. 3C); this is consistent with previous human microbiome studies that have sampled multiple sites of the human colon (21–23). Thus interpersonal variation drove the differences between samples more than whether the sample came from the proximal or distal side of the colon or from the lumen or mucosa.

### Random Forest classification models identify important OTUs on each side

To identify OTUs that were distinct at each site, we constructed several Random Forest models trained using OTU relative abundances. We built the first model to classify the luminal versus mucosal samples for the proximal and distal sides, independently (Fig. 4A). The models performed well when classifying these samples (proximal AUC = 0.716, distal AUC = 0.862). The OTUs that were most predictive of each site were identified by their greatest mean decrease in accuracy when removed from the model. For distinguishing the proximal lumen and mucosa, OTUs affiliated with the *Bacteriodes, Actinomyces, Psuedomonas* and *Enterobacteraceae* were included in the best model (Fig. 5A). The model to differentiate between the distal lumen and mucosa included OTUs affiliated with the *Turicibacter, Finegoldia, Peptoniphilus* and *Anaerococcus* (Fig. 5B). These results indicated that there were fine differences between the different sites of the colon, and that these could be traced to specific OTUs on each side.

**Figure 4.**
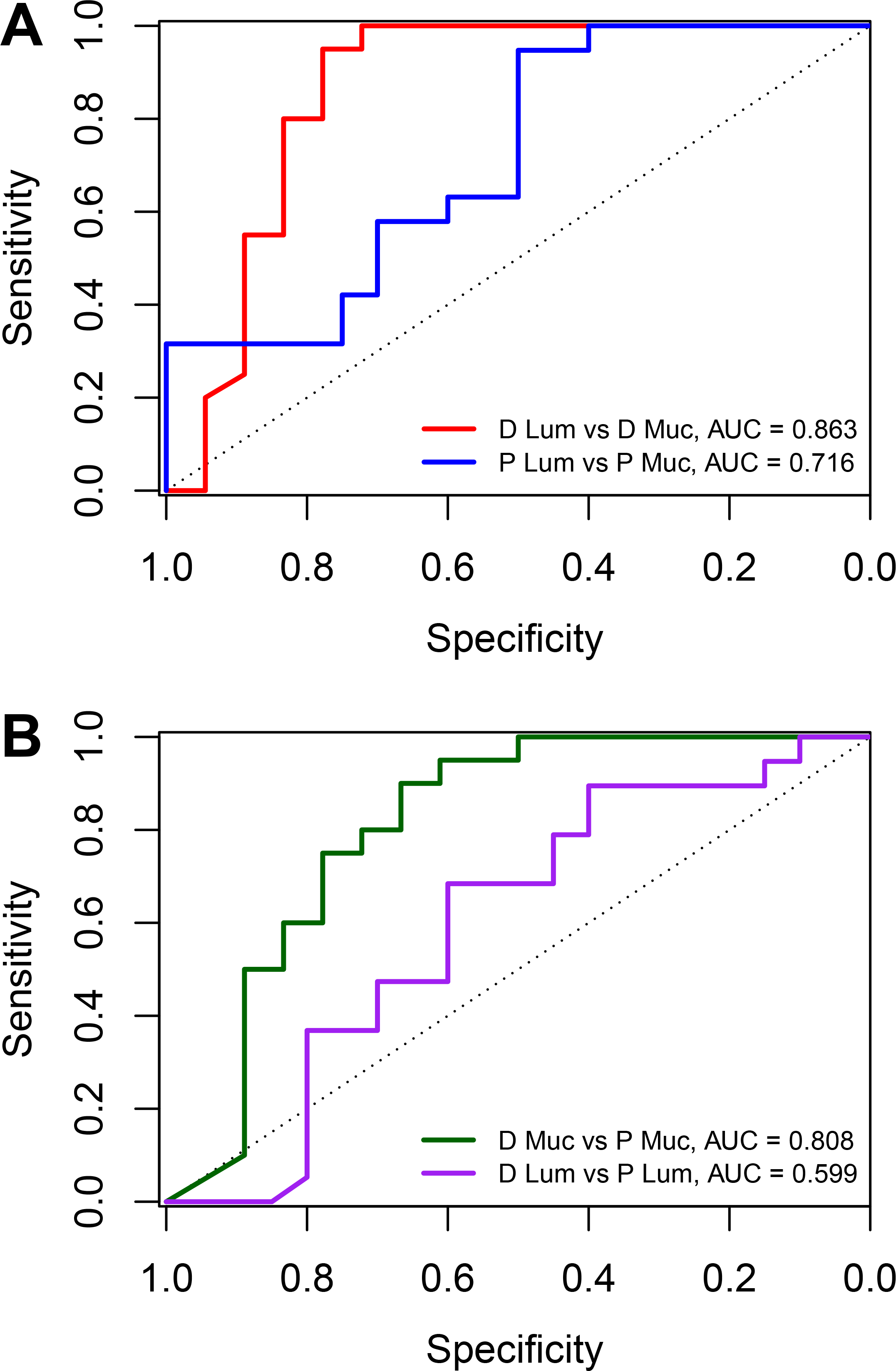
Random Forest classifies locations in the colon. A) Receiver Operator Characteristic curves are shown for the Random Forest model classifying lumen and mucosal samples for the distal (red) and proximal (blue) sides of the colon. (B) Receiver Operator Characteristic curves are shown for the 10-fold cross validation of the Random Forest model classifying distal mucosa vs proximal mucosa (green) and distal lumen versus proximal lumen (purple).

**Figure 5.**
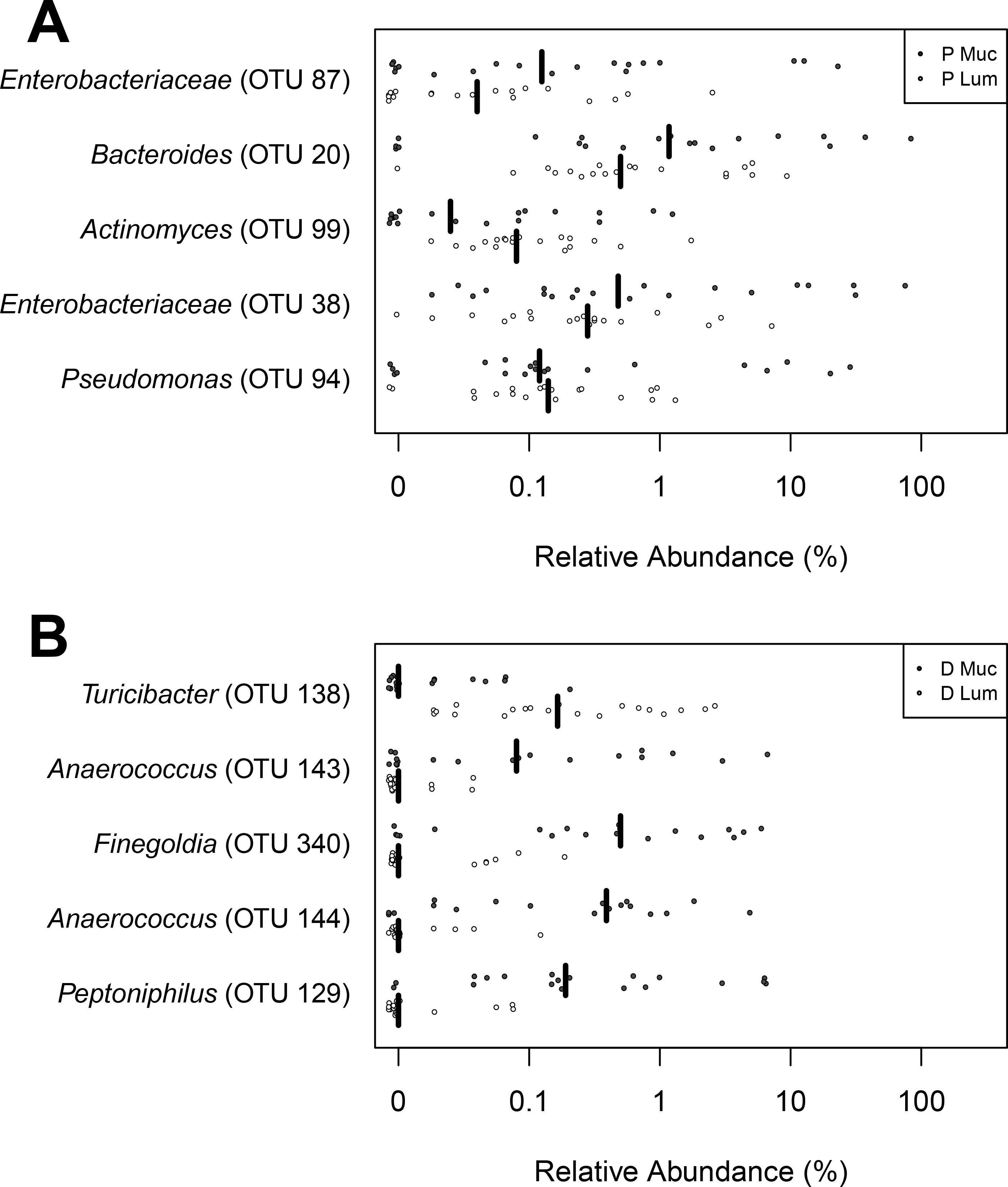
Taxa specific to the distal and proximal sides of the colon. Top five OTUs that are most important for the classification model for the distal mucosa and lumen (A) and the proximal mucosa and lumen (B). The vertical lines represent the median values for each OTU.

Next, we built a Random Forest model to differentiate the proximal and distal luminal samples. The model performed best when distinguishing the proximal versus distal mucosa (Fig. 4B, AUC = 0.808) whereas the model to differentiate between the proximal versus distal lumen performed poorly (AUC = 0.599). The OTUs included in the model differentiating the distal and proximal mucosa included members of the *Porphyromonas, Murdochiella, Finegoldia, Anaerococcus* and *Peptoniphilus* (Fig. 6A). The model that attempted to separate the the proximal and distal lumen included OTUs affiliated with the *Bacteroides, Clostridium IV* and *Oscillibacter* (Fig. 6B). Interestingly, *Anaerococcus* and *Finegoldia* were distinct between the mucosa and lumen and also helped to differentiate between the proximal and distal sides.

**Figure 6.**
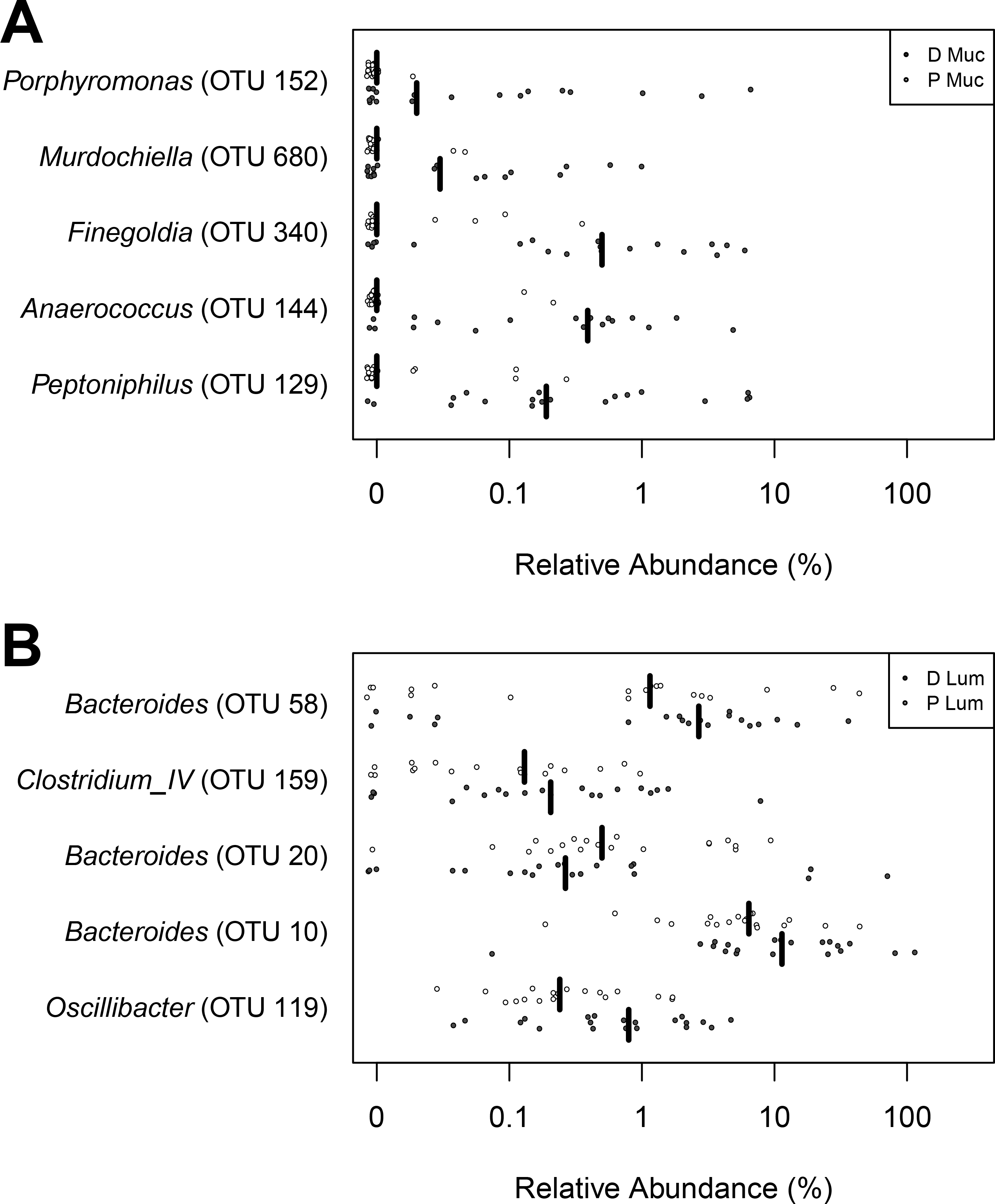
Taxa specific to the distal and proximal mucosa and lumen. The five OTUs that were most important differentiating the distal and proximal mucosa (A) and the distal and proximal lumen (B). The vertical lines represent the median values for each OTU.

### Bacterial OTUs associated with CRC and IBD are found in healthy individuals

Given that specific bacterial species have been associated with colorectal cancer and IBD, we probed our sample set for these OTUs. Among our 100 samples, the most frequent sequence associated with the *Fusobacterium* genus was OTU179, which aligned via blastn to *Fusobacterium nucleatum subsp animalis* (100% over full length). This is the only species of *Fusobacterium* known to have oncogenic properties and be found on the surfaces of colorectal cancer tumors (24). There were 14 samples from 8 subjects with the *F. nucleatum subsp. animalis* sequences. Of the samples with the highest relative abundance of *F. nucleatum subsp. animalis*, four of the samples were from the proximal mucosa and three from the distal mucosa (Supplementary Fig. S1A). The second most frequent *Fusobacterium* sequence was OTU472, which aligned with 99% identity to *F. varium*. In addition to *F. nucleatum, F. varium* has been associated with IBD (25). Four subjects harbored *F. varium* and the samples were split evenly between the proximal and distal mucosa (Supplementary Fig. S1B). OTU152 was similar to the members of the *Porphyromonas* genus and the most frequent sequence in that OTU aligned to *Porphyromonas asacharolytica* (99% over full length), another bacterium commonly detected and isolated from colorectal tumors. OTU152 was only detected on the distal mucosa, and in fact was one of the OTUs the classification model identified as separating distal and proximal sides (Supplementary Fig. S1C). Among the 11 distal mucosa samples that were positive for *P. asacharolytica*, the relative abundances for this OTU ranged from 0.01% to 16%. Thus, disease-associated OTUs could be found in our sample set of 20 healthy individuals.

## Discussion

We identified bacterial taxa that were specific to the lumen and mucosa of the proximal and distal sides of the human colon using samples collected during an unprepared colonoscopy of healthy subjects. We found that all locations contained a range of phylum relative abundances and a range of diversity, but that there was a wide variability between subjects. Pairwise comparisons of each of the sites revealed that the proximal mucosa and lumen were most similar to each other. Further, comparison of colonoscopy-collected samples with fecal samples demonstrated that the distal lumen was most similar to feces. Random Forest models built using OTU relative abundances from each sample identified microbiota that were particular to each location of the colon. Finally, we were able to detect some bacterial OTUs associated with colonic disease in our healthy cohort. Using unprepped colonoscopies and machine learning, we have identified bacterial taxa specific to the healthy proximal and distal human colon.

When examining the relative abundance of the dominant phyla at each site (i.e. *Bacteriodes* and *Firmicutes*), there was a wide amount of variation. This likely reflects not only the variability between human subjects, caused by differences in age, sex, and diet, but may also reflect spatial patchiness in the gut microbiome within a subject. Patchiness refers to inconsistent distribution of microbial populations due to fluctuations in local resources (26). One study noted that the bacteria recoverable from the same mucosal sample location can be vastly different when the samples are taken just 1 cm away from each other (27). Similar patchiness was also observed in luminal contents and fecal samples themselves; there was separation of different interacting microbes along the length of a stool sample, for instance (28). A third study that sampled six mucosal sites along the colon observed such patchiness in two of the three study subjects (21). While our subjects were not sampled frequently enough to draw specific conclusions about patchiness along the unprepped colon, we did observe some specific differences in mucosal versus luminal samples at the phylum level. The mucosal samples harbored more *Proteobacteria*, consistent with previous studies comparing mucosal swabs to luminal content in humans (4). However, we must still consider that the results from phyla analysis may have been impacted by inter-subject patchiness.

To get around the noisiness from a diverse set of samples, we built Random Forest classification models to identify the microbiota that were specific to each side and in the lumen and mucosa. For each comparison we identified the top five OTUs that were strongly predictive of one site or another. Generally, OTUs identified in each location were consistent with known physiological gradients along the gut axis (5). For instance, the proximal mucosa contains the highest oxygen concentrations of the colon and harbored mucosa-associated facultative anaerobes such as *Actinomyces* and *Enterobacteraceae* and aerobic *Psuedomonas*. The distal mucosa was far more likely to host strictly anaerobic species such as *Porphyromonas, Anaerococcus, Finegoldia* and *Peptoniphilus*. Thus the gut microenvironment of each location likely enriches for these specific microbiota.

In addition to identifying features that are specific to each side of the gut, the ability of the Random Forest to classify samples can serve as a proxy for similarity. That is, a higher AUC value indicates the samples are more efficiently classfied (and thus more different) than a model with a lower AUC value. For instance, the model separating the proximal and distal mucosa had an AUC of 0.850 whereas the model for classifying the proximal and distal lumen had a much lower AUC of 0.580. Further, the latter model required 44 OTUs to best separate the samples whereas the models separating the mucosa only needed 10 OTUs. The much lower AUC and need for a high number of features compared to other models suggest these locations are the most similar of the comparisons tested. We speculate that the model was less effective at classifying the proximal and distal luminal contents because the mucosal microenvironments have more variable selective pressure along the colon than the luminal microenvironments.

We detected *F. nucleatum* and *P. asacharolytica* in 8 and 5 of our subjects, respectively. These bacteria have been shown to be predictive of colorectal cancer in humans (9) and have oncogenic properties in cell culture and in mice (29). Though the bacteria are known to co-localize on CRC tumors, in our study *F. nucleatum* was found on both sides of the colon while *P. asacharolytica* was only detected in the distal mucosa. Not much is known about the distribution of *P. asacharolytica* along the healthy colon, but given its anaerobic lifestyle and asacharolytic metabolism, it is perhaps not surprising that our study detected the bacteria primarily in the less-oygen-rich and protein-rich distal mucosa (4). In studies examining bacteria on colorectal cancer tumors, *F. nucleatum* was more commonly detected on proximal-sided tumors, and distribution of *F. nucleatum* decreased along the colon to rectum (30). Of the 8 (40%) individuals positive for *F. nucleatum* in our study, the bacterium was spread across the proximal mucosa, distal lumen and distal mucosa. The *Fusobacterium* species *nucleatum* and *varium* have been commonly isolated from mucosal biopsies of patients with IBD and UC (25,31). In our study, *F. varium* was only detected in three subjects and two of those samples were isolated form the proximal mucosa (Supplementary Fig. S1B). *F. varium* is most commonly isolated from UC patient biopsies from the ileum or cecum (adjacent to the proximal colon) (32), suggesting this species may exhibit preference for the higher oxygen content of these gastrointestinal sites.

Spatial organization of *Fusobacteria* and other bacterial species into polymicrobial biofilms that can invade the gut mucosa have been linked to CRC (33). The biofilms promote tumorigenesis by allowing bacteria to grow near the epithelium, inducing inflammation, genotoxicity and metabolic changes that favor tumor cell growth (33). In one study of CRC biofilms and tumors, all of the proximal tumors examined contained a polymicrobial biofilm on the tumor mucosa (7). Further examination of these tumors identified *Fusobacteria* species as members of the biofilm. These results indicate that it is not only the presence of the bacterium but the tumor community as a whole that contributes to tumorigenesis (7). A ‘driver-passenger’ model has been proposed as a mechanism for biofilm assembly in the gut (34,35). In this model, ‘driver’ species such as *Fusobacterium spp* and *Porphyromonas spp* exert tumorigenic effects locally and create a niche for adherence of ‘passenger’ species that comprise the rest of the biofilm (34,35). Thus the the distribution of these disease-associated microbes in healthy patients is of interest as their presence can be predictive of disease prior to the onset of symptoms (9). A better understanding of the early microbial changes in the gut microboiome is essential for elucidating a mechanism for development of CRC or IBD subtypes in the proximal or distal colon.

Specific comparisons of our findings to previously published studies of spatial variation are confounded by the use of bowel preparation methods. A rare report of a matched-colonoscopy study sampled 18 patient’s colonic mucosa and luminal contents prior to and after bowel cleansing (36). This study found that mucosal and luminal samples were distinguishable prior to bowel cleansing, but that bowel preparation resulted in an increase in shared OTUs between each site (36). After seven days, bowel cleansing not only made the samples more difficult to distinguish, but it also decreased the diversity observed across sites. Bowel preparation clearly biases the representation of microbiota recovered from sampling the lumen or mucosa.

By revealing specific differences in microbial populations at each location in the gut via sampling an unprepared bowel, we can begin to form hypotheses about how specific host-microbe interactions can affect disease progression of proximal and distal CRC and IBD subtypes. Future investigation of these samples using metagenomics and metatranscriptomics would illuminate the microbial activities in these gut microenvironments. Further, combining this approach with a more comprehensive sampling strategy along the unprepped colon could enhance microbiome-based screening and treatment modalities for colon disease.

## Acknowledgments

We thank all the individuals who volunteered for the study. This work was supported by the Rose and Lawrence C. Page Foundation (DKT). We would also like to thank Brian Kleiner, Chelsea Crofoot, and Kirk Herman for their roles in study coordination, subject recruitment, sample collection and sample processing. Thanks also to GI Fellows Drs Amy Hosmer, Alexander Larson and Anna Tavakkoli who assisted Dr. Turgeon with the colonoscopies.

## Figure S1

Location and relative abundance of cancer-associated OTUs. Relative abundance was calculated and plotted by sample site for each OTU of interest: (A) *Fusobacterium nucleatum subsp. animalis* (B) *Fusobacterium varium* and (C) *Porphyromonas asacharolytica*

## References

1. Yamauchi M, LochheadP, MorikawaT, HuttenhowerC, ChanAT, GiovannucciE, et al. Colorectal cancer: A tale of two sides or a continuum?: Figure 1. Gut [Internet]. BMJ; 2012;61:794–7. Available from: https://doi.org/10.1136/gutjnl-2012-302014

2. ForbesJD, DomselaarGV, BernsteinCN. The gut microbiota in immune-mediated inflammatory diseases. Frontiers in Microbiology [Internet]. Frontiers Media SA; 2016;7. Available from: https://doi.org/10.3389/fmicb.2016.01081.

3. Halfvarson J, BrislawnCJ, LamendellaR, Vazquez-BaezaY, WaltersWA, BramerLM, et al. Dynamics of the human gut microbiome in inflammatory bowel disease. Nature Microbiology [Internet]. Springer Nature; 2017;2: 17004. Available from: https://doi.org/10.1038/nmicrobiol.2017.4

4. Albenberg L, EsipovaTV, JudgeCP, BittingerK, ChenJ, LaughlinA, et al. Correlation between intraluminal oxygen gradient and radial partitioning of intestinal microbiota. Gastroenterology [Internet]. Elsevier BV; 2014;147:1055–1063.e8. Available from: https://doi.org/10.1053/j.gastro.2014.07.020

5. DonaldsonGP, LeeSM, MazmanianSK. Gut biogeography of the bacterial microbiota. Nature Reviews Microbiology [Internet]. Springer Nature; 2015;14:20–32. Available from: https://doi.org/10.1038/nrmicro3552.

6. KosticAD, ChunE, RobertsonL, GlickmanJN, GalliniCA, MichaudM, et al. Fusobacterium nucleatum potentiates intestinal tumorigenesis and modulates the tumor-immune microenvironment. Cell Host & Microbe [Internet]. Elsevier BV; 2013;14:207–15. Available from: https://doi.org/10.1016%2Fj.chom.2013.07.007.

7. DejeaCM, WickEC, HechenbleiknerEM, WhiteJR, WelchJLM, RossettiBJ, et al. Microbiota organization is a distinct feature of proximal colorectal cancers. Proceedings of the National Academy of Sciences [Internet]. Proceedings of the National Academy of Sciences; 2014;111: 18321–6 Available from: https://doi.org/10.1073/pnas.1406199111

8. McCoy AN, Araújo-Pérez F, Azcárate-PerilA, YehJJ, SandlerRS, KekuTO. Fusobacterium is associated with colorectal adenomas. Goel A, editor. PLoS ONE [Internet]. Public Library of Science (PLoS); 2013;8:e53653. Available from: https://doi.org/10.1371%2Fjournal.pone.005365.

9. BaxterNT, RuffinMT, RogersMAM, SchlossPD. Microbiota-based model improves the sensitivity of fecal immunochemical test for detecting colonic lesions. Genome Medicine [Internet]. Springer Nature; 2016;8. Available from: https://doi.org/10.1186/s13073-016-0290-3.

10. Strauss J, KaplanGG, BeckPL, RiouxK, PanaccioneR, DeVinney R, et al. Invasive potential of gut mucosa-derived fusobacterium nucleatum positively correlates with IBD status of the host. Inflammatory Bowel Diseases [Internet]. Ovid Technologies (Wolters Kluwer Health); 2011;17: 1971–8. Available from: https://doi.org/10.1002/ibd.21606.

11. BrennanCA, GarrettWS. Gut microbiota, inflammation, and colorectal cancer. Annual Review of Microbiology [Internet]. Annual Reviews; 2016;70:395–411. Available from: https://doi.org/10.1146%2Fannurev-micro-102215-095513.

12. Flemer B, LynchDB, BrownJMR, JefferyIB, RyanFJ, ClaessonMJ, et al. Tumour-associated and non-tumour-associated microbiota in colorectal cancer. Gut [Internet]. BMJ Publishing Group; 2017;66:633–43. Available from: http://gut.bmj.com/content/66/4/633.

13. Jalanka J, SalonenA, Salojärvi J, RitariJ, ImmonenO, MarcianiL, et al. Effects of bowel cleansing on the intestinal microbiota. Gut [Internet]. BMJ; 2014;64:1562–8. Available from: https://doi.org/10.1136/gutjnl-2014-307240.

14. Harrell L, WangY, AntonopoulosD, YoungV, LichtensteinL, HuangY, et al. Standard colonic lavage alters the natural state of mucosal-associated microbiota in the human colon. SinghSR, editor.PLoS ONE [Internet]. Public Library of Science (PLoS); 2012;7:e32545. Available from: https://doi.org/10.1371/journal.pone.0032545

15. KozichJJ, WestcottSL, BaxterNT, HighlanderSK, SchlossPD. Development of a dual-index sequencing strategy and curation pipeline for analyzing amplicon sequence data on the MiSeq illumina sequencing platform. Applied and Environmental Microbiology [Internet]. American Society for Microbiology; 2013;79:5112–20. Available from: https://doi.org/10.1128/aem.01043-13.

16. SchlossPD, WestcottSL, RyabinT, HallJR, HartmannM, HollisterEB, et al. Introducing mothur: Open-source, platform-independent, community-supported software for describing and comparing microbial communities. Applied and Environmental Microbiology [Internet]. American Society for Microbiology; 2009;75:7537–41. Available from: https://doi.org/10.1128/aem.01541-09

17. Wang Q, GarrityGM, TiedjeJM, ColeJR. Naive bayesian classifier for rapid assignment of rRNA sequences into the new bacterial taxonomy. Applied and Environmental Microbiology [Internet]. American Society for Microbiology; 2007;73:5261–7. Available from: https://doi.org/10.1128/aem.00062-07.

18. CalleML, UrreaV, Boulesteix A-L, MalatsN. AUC-RF: A new strategy for genomic profiling with random forest. Human Heredity [Internet]. S. Karger AG; 2011;72:121–32. Available from: https://doi.org/10.1159%2F000330778.

19. Robin X, TurckN, HainardA, TibertiN, LisacekF, Sanchez J-C, et al. pROC: An open-source package for r and s to analyze and compare ROC curves. BMC Bioinformatics [Internet]. Springer Nature; 2011;12:77. Available from: https://doi.org/10.1186%2F1471-2105-12-77.

20. Lloyd-Price J, Abu-AliG, HuttenhowerC. The healthy human microbiome. Genome Medicine [Internet]. Springer Nature; 2016;8. Available from: https://doi.org/10.1186/s13073-016-0307-y.

21. Eckburg PB. Diversity of the human intestinal microbial flora. Science [Internet]. American Association for the Advancement of Science (AAAS); 2005;308:1635–8. Available from: https://doi.org/10.1126/science.1110591

22. Cárcer DA de, CuívPÓ, WangT, KangS, WorthleyD, WhitehallV, et al. Numerical ecology validates a biogeographical distribution and gender-based effect on mucosa-associated bacteria along the human colon. The ISME Journal [Internet]. Springer Nature; 2010;5:801–9. Available from: https://doi.org/10.1038/ismej.2010.177.

23. Zhang Z, GengJ, TangX, FanH, XuJ, WenX, et al. Spatial heterogeneity and co-occurrence patterns of human mucosal-associated intestinal microbiota. The ISME Journal [Internet]. Springer Nature; 2013;8:881–93. Available from: https://doi.org/10.1038/ismej.2013.185.

24. Castellarin M, WarrenRL, FreemanJD, DreoliniL, KrzywinskiM, StraussJ, et al. Fusobacterium nucleatum infection is prevalent in human colorectal carcinoma. Genome Research [Internet]. Cold Spring Harbor Laboratory Press; 2011;22:299–306. Available from: https://doi.org/10.1101/gr.126516.111.

25. Lee Y, EunCS, LeeAR, ParkCH, HanDS. FusobacteriumIsolates recovered from colonic biopsies of inflammatory bowel disease patients in korea. Annals of Laboratory Medicine [Internet]. Korean Society for Laboratory Medicine (KAMJE); 2016;36:387. Available from: https://doi.org/10.3343/alm.2016.36.4.387.

26. Rougharden JD. Patchiness in the spatial distribution of a population caused by stochastic fluctuations in resources. Oikos [Internet]. JSTOR; 1977;29:52. Available from: https://doi.org/10.2307/3543292.

27. Hong P-Y, CroixJA, GreenbergE, GaskinsHR, MackieRI. Pyrosequencing-based analysis of the mucosal microbiota in healthy individuals reveals ubiquitous bacterial groups and microheterogeneity. Ahmed N, editor. PLoS ONE [Internet]. Public Library of Science (PLoS); 2011;6:e25042. Available from: https://doi.org/10.1371/journal.pone.0025042.

28. StearnsJC, LynchMDJ, SenadheeraDB, TenenbaumHC, GoldbergMB, CvitkovitchDG, et al. Bacterial biogeography of the human digestive tract. Scientific Reports [Internet]. Springer Nature; 2011;1. Available from: https://doi.org/10.1038/srep00170

29. SearsCL, GarrettWS. Microbes, microbiota, and colon cancer. Cell Host & Microbe [Internet]. Elsevier BV; 2014;15:317–28. Available from: https://doi.org/10.1016/j.chom.2014.02.007.

30. Mima K, CaoY, ChanAT, QianZR, NowakJA, MasugiY, et al. Fusobacterium nucleatum in colorectal carcinoma tissue according to tumor location. Clinical and Translational Gastroenterology [Internet]. Springer Nature; 2016;7:e200. Available from: https://doi.org/10.1038/ctg.2016.53.

31. Ohkusa T. Induction of experimental ulcerative colitis by fusobacterium varium isolated from colonic mucosa of patients with ulcerative colitis. Gut [Internet]. BMJ; 2003;52:79–83. Available from: https://doi.org/10.1136/gut.52.1.79.

32. Ohkusa T, SatoN, OgiharaT, MoritaK, OgawaM, OkayasuI. Fusobacterium varium localized in the colonic mucosa of patients with ulcerative colitis stimulates species-specific antibody. Journal of Gastroenterology and Hepatology [Internet]. Wiley-Blackwell; 2002;17:849–53. Available from: https://doi.org/10.1046/j.1440-1746.2002.02834.x.

33. Li S, KonstantinovSR, SmitsR, PeppelenboschMP. Bacterial biofilms in colorectal cancer initiation and progression. Trends in Molecular Medicine [Internet]. Elsevier BV; 2017;23:18–30. Available from: https://doi.org/10.1016/j.molmed.2016.11.004.

34. Tjalsma H, BoleijA, MarchesiJR, DutilhBE. A bacterial driverpassenger model for colorectal cancer: Beyond the usual suspects. Nature Reviews Microbiology [Internet]. Springer Nature; 2012;10:575–82. Available from: https://doi.org/10.1038/nrmicro281.

35. FlynnKJ, BaxterNT, SchlossPD. Metabolic and community synergy of oral bacteria in colorectal cancer. McMahon K, editor. mSphere [Internet]. American Society for Microbiology; 2016;1:e00102–16. Available from: https://doi.org/10.1128/msphere.00102-16.

36. ShobarRM, VelineniS, KeshavarzianA, SwansonG, DeMeoMT, MelsonJE, et al. The effects of bowel preparation on microbiota-related metrics differ in health and in inflammatory bowel disease and for the mucosal and luminal microbiota compartments. Clinical and Translational Gastroenterology [Internet]. Springer Nature; 2016;7:e143. Available from: https://doi.org/10.1038/ctg.2015.54.

